# READemption – A tool for the computational analysis of deep-sequencing-based transcriptome data

**DOI:** 10.1101/003723

**Authors:** Konrad U. Förstner, Jörg Vogel, Cynthia M. Sharma

## Abstract

**Summary:** RNA-Seq has become a potent and widely used method to qualitatively and quantitatively study transcriptomes. In order to draw biological conclusions based on RNA-Seq data, several steps some of which are computationally intensive, have to be taken. Our READemption pipeline takes care of these individual tasks and integrates them into an easy-to-use tool with a command line interface. To leverage the full power of modern computers, most subcommands of READemption offer parallel data processing. While READemption was mainly developed for the analysis of bacterial primary transcriptomes, we have successfully applied it to analyze RNA-Seq reads from other sample types, including whole transcriptomes, RNA immunoprecipitated with proteins, not only from bacteria, but also from eukaryotes and archaea.

**Availability and Implementation:** READemption is implemented in Python and is published under the ISC open source license. The tool and documentation is hosted at http://pythonhosted.org/READemption (DOI:10.6084/m9.figshare.977849).

## 1 INTRODUCTION

RNA-Seq, the examination of cDNA by massively parallel sequencing technologies, is a potent way to perform transcriptome analyses at single-nucleotide-resolution and with a high dynamic range (Wang *et al.*, 2009). It has been successfully used to annotate transcript boundaries and to identify novel transcripts such as small regulatory RNAs in both pro- and eukaryotes (Filiatrault, 2011; Ozsolak and Milos, 2011). Most prominently, it can be applied to quantify the expression levels of genes and is about to replace microarrays. It can also be used to study the interaction of proteins and RNA through performing RNA-Seq of co-immunoprecipation (coIP) samples (König *et al.*, 2012). Likewise, any other subset of RNA molecules from, for instance, RNA size-fractionation, ribosome profiling, metatranscriptomics, or degradome profiling experiments can be sequenced. Due to ever lower cost and increasing speed of sequencing, the bioinformatical analysis has become a bottleneck of RNA-Seq-based projects.

We have created an automated RNA-Seq processing pipeline named READemption with the initial purpose to handle differential RNA-Seq (dRNA-Seq) data for the determination of transcriptional start sites in bacteria (Sharma *et al.*, 2010). We saw the need for this as other available RNA-Seq analysis pipelines (e.g. Delhomme *et al.*, 2012, McClure *et al.*, 2013) were not designed for this application. Additionally, while most available RNA-Seq pipelines put priority on fast mapping we have chosen a short read aligner with a relatively high demand of memory and computation capacities as in return it offers high sensitivity as well as a low false-discovery rate and can perform multiple splitting of reads (Hoffmann *et al.*, 2009). We have since extended the functionality of this Python-based pipeline so it is now capable of analyzing RNA-Seq reads from a wide range of experiments. We have successfully applied READemption for the analyses of RNA samples from bacterial, archaeal and eukaryotic species as well as for RNA virus genomes (e.g. Dugar *et al.*, 2013; Zhelyazkova *et al.*, 2012). It is able to work with reads from both Illumina and 454 platforms of different lengths and can be used for single- and paired-end sequenced libraries.

## 2 DESCRIPTION

READemption integrates the steps that are required to interpret and gain biological knowledge from RNA-Seq experiments in one tool and makes them accessible via a consistent command line interface. Additionally, it conducts parallel data processing to reduce the run time. The tool performs clipping and size filtering of raw cDNA reads from different sequencing platforms, mapping to reference sequences, coverage calculation, gene-based quantification and comparison of expression levels. Moreover, it provides several statistics (e.g. of the read mappability) and generates plots as well as files for visualization of the results in genome browsers. READemption was designed as high-performance application and follows the concept of “convention over configuration”. This includes the use of established default parameter values and the approach that files are placed or linked into defined paths and are then treated accordingly. Though this design principle, READemption offers several parameters which enable the user to adapt its execution to the specific needs.

READemption provides the subcommands: `align`, `coverage`, `gene_quanti`, `deseq`, `viz_align`, `viz_gene_quanti` and `viz_deseq` which combine several processing substeps into comprehensible units.

*Read processing and mapping*: The fundamental tasks of pre-processing the input reads and aligning them to reference sequences is covered by the subcommand `align`. In an initial step READemption removes poly(A)-tails introduced during the library preparation and discards too short reads. For the alignment of reads to reference sequences, the short read mapper `segemehl` and its remapper `lack` (Hoffmann *et al.*, 2009) are used. The mapping is followed by the conversion of the resulting SAM alignment files into BAM files and the generation of mapping statistics.

*Coverage calculation*: Based on the alignments provided in the BAM files, cDNA coverage files can be generated using the subcommand `coverage`. It creates several wiggle files that are based on different normalization methods like total number of aligned reads and represent the nucleotide-wise cDNA coverage in a strand specific manner. In order to visually inspect the libraries, these wiggle files can be loaded into common genome browsers.

**Fig. 1.**
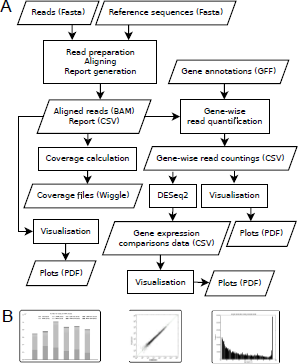
A) Data and work flow of READemption including the input, output and the performed steps. B) Examples of plots generated by READemption

*Gene expression quantification*: The read alignments can also be further used by the subcommand `gene_quanti` to calculate gene-wise read counts. For this purpose, annotation files including gene positions in GFF3 format have to be provided. Besides raw gene-wise read countings normalized values – by total number of aligned reads as well as RPKM (Mortazavi *et al.*, 2008) – are returned.

*Differential gene expression analysis*: For pairwise expression comparison, the subcommand `deseq` offers statistical analysis based on the approach implemented in `DESeq2` (Anders and Huber, 2010) which builds upon the raw read counting. The results of `DESeq2` are reformated and supplemented with gene annotations.

*Plotting*: The final three subcommands called `viz_align`, `viz_genequanti` and `viz_deseq` generate several visualizations that help to interpret the result of the subcommand `align`, `gene_quanti` and `deseq`, respectively.

READemption requires `Python` 3.2 or higher (http://python.org) and the libraries `matplotlib` (Hunter, 2007) and `numpy` (Oliphant, 2007) as well as the `samtools` (Li *et al.*, 2009) wrapper `pysam` (http://pypi.python.org/pypi/pysam/) are needed. The subcommand `deseq` relies on an `R` (http://cran.r-project.org) installation and the bioconductor package `DESeq2` (Anders and Huber, 2010). Instructions how to install READemption as well as how to execute its subcommands including examples can be found in the documentation.

## 3 CONCLUSIONS

We present an open source pipeline for the analysis of RNA-Seq data from all domains of life. READemption generates several output files that can be examined with common office suites, graphic programs and genome browsers. Its features make it a useful tool for anybody interested in the computational analysis of RNA-Seq data with the required basic command line skills.

## ACKNOWLEDGEMENT

We thank members of the Sharma and the Vogel groups especially Thorsten Bischler for testing and constructive feedback.

*Funding*: Work in the Sharma laboratory is supported by the ZINF Young Investigator program of the Research Center for Infectious Diseases (ZINF) at the University of Würzburg and the Bavarian BioSysNet program. CMS is supported by the Young Academy program of the Bavarian Academy of Sciences.

